# Dose- and time-dependent effects of cobalt protoporphyrin IX on granulocyte mobilization and metabolic markers in mice

**DOI:** 10.1101/2024.10.22.619673

**Authors:** Aleksandra Bednarz, Paweł Kozuch, Kacper Kowalski, Izabella Skulimowska, Neli Kachamakova-Trojanowska, Jadwiga Filipek-Gorzała, Patrycja Kwiecinska, Raquel Garcia Garcia, Kinga Gawlinska, Kinga Malezyna, Andrzej Kubiak, Natalia Bryniarska-Kubiak, Alicja Jozkowicz, Krzysztof Szade, Agata Szade

## Abstract

Recombinant granulocyte colony-stimulating factor (G-CSF) is the most commonly used agent for treating neutropenia and mobilizing hematopoietic stem cells (HSCs) for transplantation. However, some patients do not respond effectively to existing mobilization protocols. To address this, the development of new therapeutic approaches is necessary. One potential strategy is the pharmacological induction of endogenous mobilizing factors, which can be achieved through the administration of cobalt protoporphyrin IX (CoPP). CoPP induces mobilization of HSCs and granulocytes by increasing endogenous G-CSF production, though the optimal dosing and potential side effects remain unclear.

The aim of our study was to optimize the dose and timing of CoPP administration and evaluate its safety in mobilizing cells from the bone marrow to the blood. Our results show that CoPP exerts a dose-dependent mobilizing effect, with the highest G-CSF levels and number of mobilized leukocytes observed in mice treated with 10 mg/kg of CoPP. While there were no severe adverse effects, there were mild fluctuations in markers of liver and kidney function, including a slight reduction in urea nitrogen (BUN) and glucose levels during the five days of administration. Additionally, although most parameters normalized within 30 days after treatment, the decrease in BUN persisted. Mice experienced short-term weight loss following CoPP administration, but they regained their initial weight within two weeks. By day 30, leukocyte counts, hematopoietic stem and progenitor cells (HSPCs) in bone marrow, and G-CSF concentration in the blood had returned to baseline.

This study demonstrates that CoPP mobilizes cells from the bone marrow to the blood in a dose-dependent manner, with mild side effects, including temporary changes in biochemical markers and a sustained reduction in BUN levels.

## Introduction

Cobalt protoporphyrin IX (CoPP) is a compound that has been widely used to induce expression of heme oxygenase-1 (HO-1 or Hmox1) since the 1970s [1]. HO-1 is an enzyme that degrades heme into ferrous ions (Fe²⁺), carbon monoxide (CO), and biliverdin, which is subsequently converted to bilirubin by biliverdin reductase [2]. The products of HO-1 activity exert various effects, including immunomodulatory [3–5], anti-apoptotic [6], and antioxidant [7] functions.

Although CoPP is commonly used as a model activator of HO-1 expression, several HO-1-independent activities of CoPP have been identified. These include the inhibition of caspase-3 and -8 activity [8], suppression of LPS-induced activation of inducible nitric oxide synthase (iNOS) in RAW264.7 macrophages [9] and upregulation of cyclooxygenase-2 (COX-2) [10].

We demonstrated that CoPP can mobilize cells from the bone marrow to the blood independently of HO-1 activity [11]. Mobilization refers to the accelerated release of hematopoietic cells from their niche in the bone marrow into peripheral blood [12,13]. This process can occur naturally, for example during infection, but can also be induced pharmacologically. Pharmacological mobilization is primarily used to treat neutropenia (an abnormally low number of granulocytes in the blood) [14] or to collect HSCs for transplantation [15,16]. The most commonly used mobilizing agent is recombinant granulocyte colony-stimulating factor (G-CSF) [17]. However, as not all patients respond effectively to currently available drugs [18], new therapeutic approaches are being explored, including the induction of endogenous mobilizing cytokines [19].

Mobilization with CoPP offers several advantages compared to recombinant G-CSF. Firstly, CoPP mobilizes a higher number of hematopoietic stem and progenitor cells (HSPCs) than recombinant G-CSF [11], which is critical for the success of hematopoietic cell transplantation (HCT) [20]. Secondly, granulocytes mobilized by CoPP exhibit a more mature phenotype, characterized by increased granularity and higher expression of the Ly6G marker, compared to those mobilized by G-CSF [11]. This is particularly significant for treating neutropenia caused by anticancer therapy, as granulocytes mobilized by G-CSF tend to be less mature and are impaired in their ability to combat certain pathogens, such as *Candida albicans* [21]. Finally, CoPP mobilizes fewer T cells than G-CSF [11]. Since T cells are the primary drivers of graft-versus-host disease, this observation might have implications for reducing the risk of this serious complication in hematopoietic cell transplantation [22]. However, further studies are needed to confirm this potential benefit.

Given those advantages of CoPP in mobilizing cells from the bone marrow to the blood, we sought to evaluate its potential as a pharmacological mobilizing agent in a mouse model. Specifically, we aimed to determine the minimal effective dose and treatment duration to minimize potential side effects. Previous study by Muhoberac et al. [23]. reported toxic effects of CoPP, evaluating its treatment as a model for cytochrome P450-centered hepatic dysfunction. The study demonstrated that CoPP causes a dose-dependent depletion of liver cytochrome P450 in rats. This finding has significant clinical implications, as cytochrome P450 enzymes play a crucial role in the metabolism of various endogenous and exogenous compounds. Disruption of these enzymes may result in altered therapeutic efficacy or increased toxicity [24,25]. Additionally, CoPP was shown to reduce the activities of NADPH-cytochrome P450 reductase and NADH-cytochrome b5 [23].

Thus, we also monitored toxicity markers during and after successful mobilization. Additionally, we assessed the long-term effects of CoPP on hematopoiesis and the function of major organs. Since CoPP has been proposed as a potential treatment for ischemic diseases [26] or diabetic wounds [27], it is crucial to fully understand its impact on the hematopoietic system and establish a treatment regimen that does not adversely affect hematopoiesis.

## Materials and Methods

### Mice

Animal work was done in accordance with the good animal practice and approved by the Local Ethical Committee for Animal Research. All experiments were performed on male C57BL/6J mice, which were bred and maintained at the Animal Facility of the Faculty of Biochemistry, Biophysics and Biotechnology. Experiments were performed in specific pathogen-free (SPF) conditions, with constant light/dark cycle (14/10h) and continuous temperature and humidity monitoring. Mice were kept in groups of ≤5 in individually ventilated cages with food and water ad libitum. Animals in each cage were randomly assigned to all experimental groups.

### Mobilization experiments

CoPP (Frontier Scientific) was dissolved in DMSO (Sigma-Aldrich) at a concentration of 200 mg/ml. The stock solution was diluted 115× in 0.9% NaCl solution. Mice were injected intraperitoneally (i.p.) at the dose of 10 mg/kg body weight (15 µmol/kg b.w.) unless stated otherwise. DMSO diluted 115× in 0.9% NaCl was used as a solvent control. Recombinant human G-CSF (rhG-CSF, Neupogen, Amgen) was used at a dose of 250 µg/kg b.w. Compounds were injected i.p. according to schedules described for each experiment. The 5-day treatment regimen, used in the majority of experiments, was chosen because it aligns with the most common clinical protocol for recombinant G-CSF in the treatment of neutropenia and hematopoietic stem cell (HSC) donor mobilization [28]. Since CoPP induces endogenous G-CSF production, this established regimen was adopted. To evaluate long-term effects, a 30-day follow-up timepoint was selected, reflecting the standard clinical evaluation period for stem cell donors post-donation, during which majority of blood parameters typically return to their normal range [28]. Unless otherwise noted, the mice were sacrificed 6 hours after the last injection. Peripheral blood (PB) was collected from vena cava at time points indicated in the description of each experiment. Complete blood count was done using Vet abc Plus+ analyzer (Horiba). Plasma samples were frozen for further analysis.

Five mice were used in each experimental group. The long-term effect of CoPP was assessed in 2 independent experiments.

### Flow cytometry

PB was depleted of erythrocytes by a hypotonic solution, washed, and stained in PBS 2% FBS for 30 minutes at 4°C. Samples were collected using LSR Fortessa cytometer (BD) and analyzed using FlowJo software (BD). The following flow cytometry reagents were used: 7AAD (BD), APC-eFluor780 rat anti-mouse CD117 (c-Kit, clone 2B8, eBioscience), PE-Cy7 rat anti-mouse Ly-6A/E (Sca-1, clone D7, BD), Alexa Fluor 700 anti-mouse Lineage Cocktail (anti-mouse CD3, clone 17A2; anti-mouse Ly-6G/Ly-6C, clone RB6-8C5; anti-mouse CD11b, clone M1/70; anti-mouse CD45R/B220, clone RA3-6B2; anti-mouse TER-119, clone Ter-119, Biolegend).

### Clinical biochemistry tests and cytokine concentration analysis

Organ toxicity markers were measured in previously frozen plasma samples using SpotChem EZ Chemistry Analyzer according to manufacturer’s instructions. The same plasma samples were also used for cytokine secretion analysis through custom designed Luminex assay kits (R&D, ThermoFisher). Signal readouts were detected with Luminex 200 system (Millipore).

### Statistical analysis

Statistical analysis was done with GraphPad Prism software. Data are presented as mean with SEM or mean with individual values. One-way ANOVA or two-way ANOVA with Tukey’s, Sidak’s or Dunnett’s posthoc tests were applied when three or more groups were compared. Grubbs’ test was used to determine significant outliers. Results were considered as statistically significant, when p ≤ 0.05 (* - p ≤ 0.05; ** - p ≤ 0.01; *** - p ≤ 0.001; **** - p ≤ 0.0001).

## Results

### CoPP-induced mobilization is dose-dependent

In our previous work, we demonstrated that a daily intraperitoneal injection of 10 mg/kg of CoPP for five days effectively mobilized cells from the bone marrow to the blood. However, in animal studies, CoPP doses range from as low as 1 mg/kg [29] to as high as 20 mg/kg [30]. As higher doses of CoPP can lead to toxicity, it is essential to explore safer dosing regimens. Therefore, we aimed to determine whether efficient mobilization could be achieved with lower doses of CoPP. We compared our standard 10 mg/kg dose of CoPP with lower doses (5 mg/kg and 1 mg/kg) in mice treated once with CoPP or a DMSO solvent control (Fig. 1a).

**Fig. 1.**
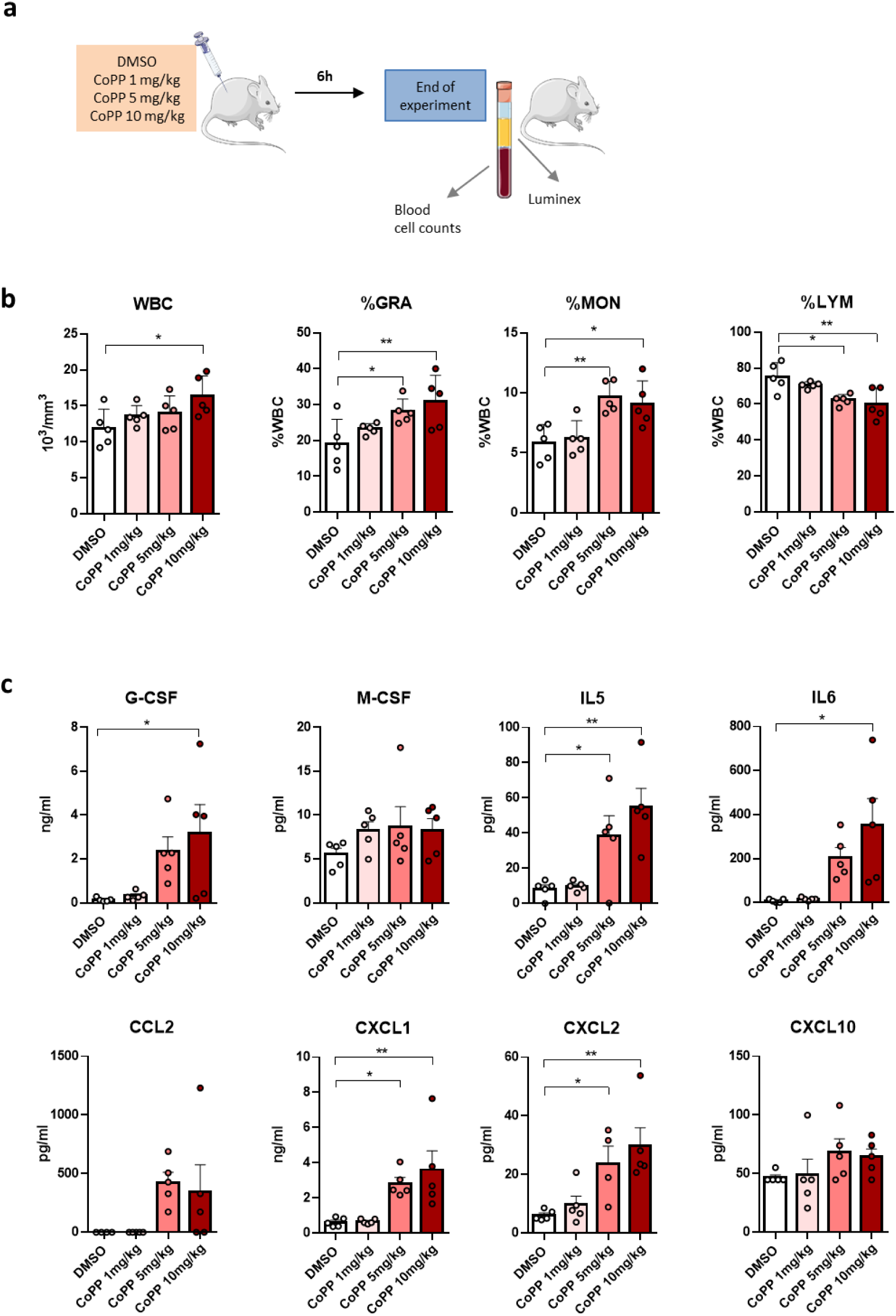
CoPP increases cytokine concentrations and induces granulocyte mobilization from the bone marrow to the blood in a dose-dependent manner. **a** Experimental scheme. C57BL/6J mice were injected once with various doses of CoPP or a solvent control (DMSO). Samples were collected 6 hours after the injection. **b** Total leukocyte count in the blood. CoPP dose-dependently increases the number of white blood cells (WBC). The percentages of granulocytes (%GRA) and monocytes (%MON) increase, while the percentage of lymphocytes (%LYM) decreases with increasing CoPP doses. **c** Selected cytokine and growth factor concentrations in plasma, measured by Luminex assay. CoPP dose-dependently increases the concentrations of G-CSF, IL-5, IL-6, CCL2 (MCP-1), CXCL1, and CXCL2. Data information: Results are shown as mean + SEM. One-way ANOVA with Dunnett’s post-hoc test, n = 5 mice per group. *p ≤ 0.05; **p ≤ 0.01; ***p ≤ 0.001; ****p ≤ 0.0001

We observed the highest number of white blood cells (WBC) in the blood of mice treated with 10 mg/kg dose of CoPP – 16 340 ±1 263 cells/µl (Fig. 1B). WBC counts in mice treated with 5 mg/kg and 1 mg/kg were lower, at 14 020 ± 1 064 and 13 600 ± 651 cells/µl, respectively, though still higher than in DMSO-treated mice (11 900 ± 1 174 cells/µl). In DMSO-treated mice, granulocytes comprised 19.02 ± 3% of WBCs, whereas with increasing doses of CoPP, the proportion of granulocytes rose to 23.22 ± 0.67% (1 mg/kg), 28.1 ± 1.56% (5 mg/kg), and 30.8 ± 3.3% (10 mg/kg) (Fig. 1b). Similarly, we observed an increase in the percentage of monocytes in the two higher-dose groups, from 5.86 ± 0.67% in DMSO-treated mice and 6.22 ± 0.65% in the 1 mg/kg group, to 9.1 ± 0.86% and 9.64 ± 0.59% in the 10 mg/kg and 5 mg/kg groups, respectively (Fig. 1b). Conversely, the percentage of lymphocytes decreased with increasing doses of CoPP, from 75.12 ± 3.5% in DMSO-treated mice to 60 ± 3.89% in the 10 mg/kg group (Fig. 1b).

The most important mediator of CoPP-induced mobilization is G-CSF. We observed increased concentrations of endogenous G-CSF in mice treated with all doses of CoPP (Fig. 1c). Even the lowest dose of 1 mg/kg led to a 2.2-fold increase in G-CSF concentration (323.6 ± 80.87 pg/ml) compared to the baseline in DMSO-treated mice (144.5 ± 39.14 pg/ml). Mice treated with 5 mg/kg showed a 16.3-fold increase (2 352 ± 650.8 pg/ml), while the highest dose of 10 mg/kg led to a 22-fold increase (3 173 ± 1 310 pg/ml) (Fig. 1c). IL-6 levels increased similarly, with a 44-fold rise observed in the 10 mg/kg group (353.2 ± 119.7 pg/ml vs. 7.98 ± 3.72 pg/ml in DMSO-treated mice). Other mobilizing cytokines, including IL-5, CCL, CXCL1, and CXCL2, were only elevated by the two higher doses (5 and 10 mg/kg), while the lowest dose (1 mg/kg) did not significantly increase their concentrations (Fig. 1c).

In summary, we found that CoPP-induced mobilization of cells from the bone marrow to the blood is dose-dependent. The minimal effective dose for achieving efficient mobilization in mice appears to be between 5 and 10 mg/kg.

### CoPP does not induce acute toxicity during a five-day treatment

The standard protocol for G-CSF-induced mobilization of granulocytes, commonly used to prevent and/or treat neutropenia associated with cancer treatment, typically requires five days of administration. In our previous studies, we followed this standard protocol to evaluate the mobilizing potential of CoPP. However, we sought to determine whether a shorter treatment duration could still achieve sufficient granulocyte mobilization. To investigate this, we conducted a time-course experiment, treating mice with CoPP, recombinant G-CSF, or control solvents daily for one to five days (Fig. 2a). We found that the highest numbers of total leukocytes and granulocytes were observed on day 5 in both groups treated with mobilizing agents (Fig. 2b). In G-CSF-treated mice, the mean leukocyte count was 23 340 ± 3 100 cells/µl, including 10 370 ± 1 770 granulocytes/µl, while in the CoPP-treated group, the mean leukocyte count was 19 980 ± 2 031 cells/µl, including 9 560 ± 1 180 granulocytes/µl. Mobilization treatments did not significantly affect erythrocyte parameters, such as hematocrit values, which ranged from 46.38 ± 2.4 (CoPP, day 4) to 53.62 ± 3.5 (CoPP, day 5, Fig. 2b).

**Fig. 2.**
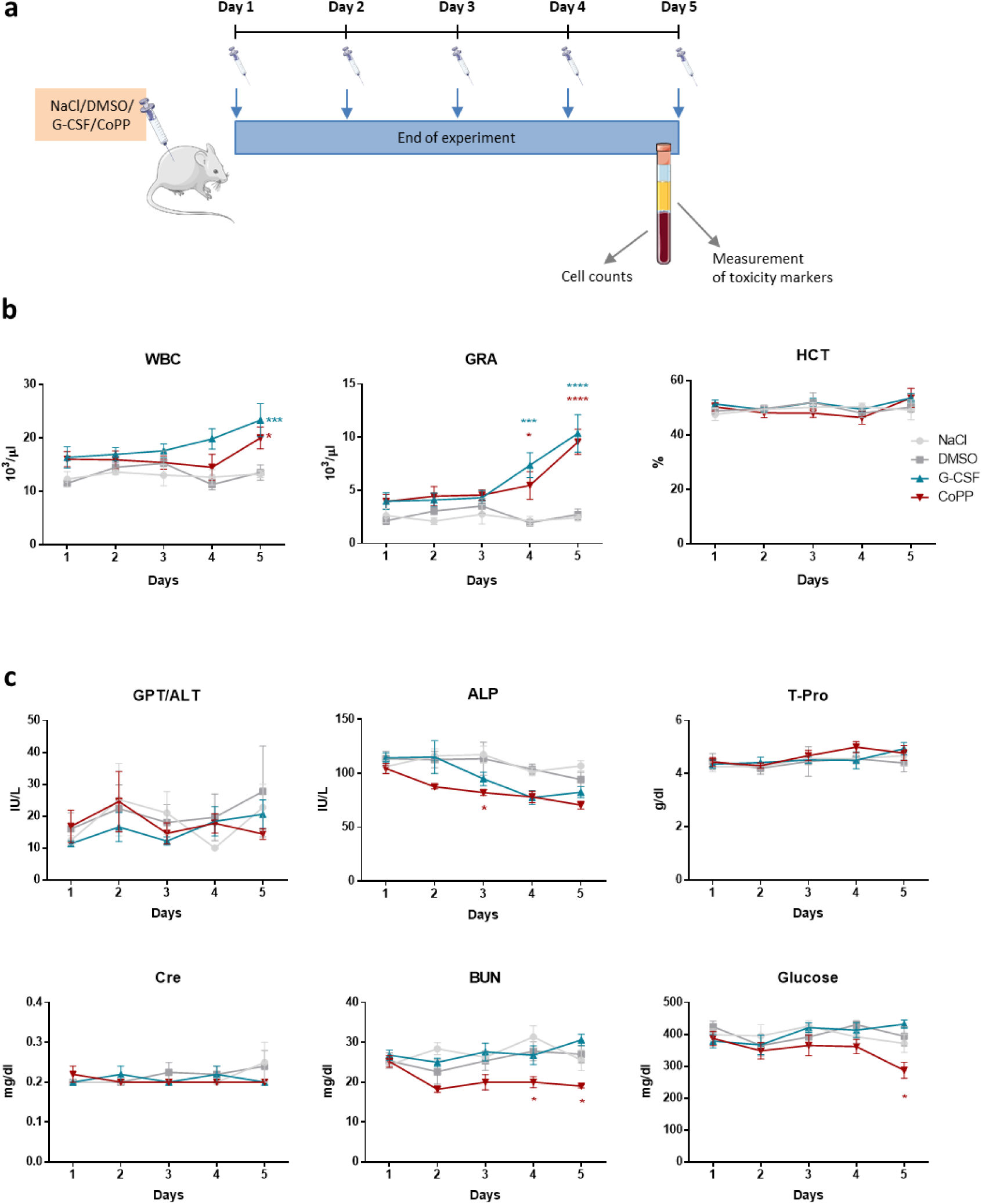
CoPP-induced mobilization of granulocytes increases with treatment duration and is associated with a decrease in plasma metabolic markers. **a** Experimental scheme. C57BL/6J mice were injected daily with 10 mg/kg CoPP, 250 µg/kg recombinant G-CSF, or solvent controls (NaCl, DMSO) for 1 to 5 days. Samples were collected 6 hours after the final injection. **b** Total leukocyte count in blood. Both recombinant G-CSF and CoPP increase the number of white blood cells (WBC) over 5 days of treatment. Granulocyte counts increase after treatment with both G-CSF and CoPP on days 1 to 3, with the greatest increases observed on days 4 and 5. A 5-day treatment with G-CSF or CoPP does not affect red blood cell parameters, such as hematocrit (HCT). **c** Plasma organ function biomarkers measured using the SpotChem EZ Chemistry Analyzer. Treatment with G-CSF or CoPP does not affect alanine transaminase (ALT/GLP) activity, total protein (T-Pro), or creatinine (Cre) levels. Both G-CSF and CoPP reduce plasma alkaline phosphatase (ALP) activity, but only CoPP, not G-CSF, decreases blood urea nitrogen (BUN) and glucose concentrations. Data information: Results are shown as mean + SEM; Two-way ANOVA with Tukey’s post-hoc test; n = 5 mice per group; *p ≤ 0.05; **p ≤ 0.01; ***p ≤ 0.001; ****p ≤ 0.0001

Previous studies have reported possible liver toxicity associated with CoPP [23]. To assess the safety of CoPP treatment in inducing mobilization, we measured biomarkers of organ function. Over the five-day treatment period with both CoPP and G-CSF, we did not observe any clear effects on alanine transaminase (GTP/ALT) activity, creatinine, or total protein concentrations (Fig. 2c). Both CoPP and G-CSF caused a slight decrease in alkaline phosphatase (ALP) activity, with the lowest levels observed on day 4 of G-CSF treatment (77.4 ± 6.46 IU/l) and day 5 of CoPP treatment (70.4 ± 3.63 IU/l), compared to control mice on day 1 (106.2 ± 5.68 IU/l for NaCl and 113.6 ± 6.74 IU/l for DMSO-treated mice). In CoPP-treated mice, we observed a decrease in blood urea nitrogen (BUN) levels starting on day 2 (18.2 ± 0.8 mg/dl, down from 25.2 ± 1.46 mg/dl on day 1) and a reduction in glucose levels, most notably on day 5 (287.4 ± 25.1 mg/dl, compared to 394.2 ± 30.37 mg/dl in DMSO-treated mice).

In summary, efficient mobilization with CoPP requires five days of continuous treatment. Importantly, this five-day CoPP regimen does not significantly induce markers of acute stress but does reduce BUN and glucose levels.

### Most of CoPP-affected parameters go back to the normal levels by day 30

Next, we aimed to evaluate the long-term effects of CoPP-induced mobilization on both the hematopoietic system and overall homeostasis in mice. Mice were treated with CoPP or G-CSF for five days, and then followed for an additional 25 days (Fig. 3a). In line with previous studies, we observed a temporary decrease in body weight in CoPP-treated mice, with the most significant reduction occurring on day 7 in the first experiment (7.1 ± 2.2% of initial weight) and on day 5 in the second experiment (2.8 ± 0.9% of initial weight). This weight loss was transient, and the mice regained their initial weight within five days after treatment ended, matching the weight of mice in other groups over the subsequent two weeks (Fig. 3b).

**Fig. 3.**
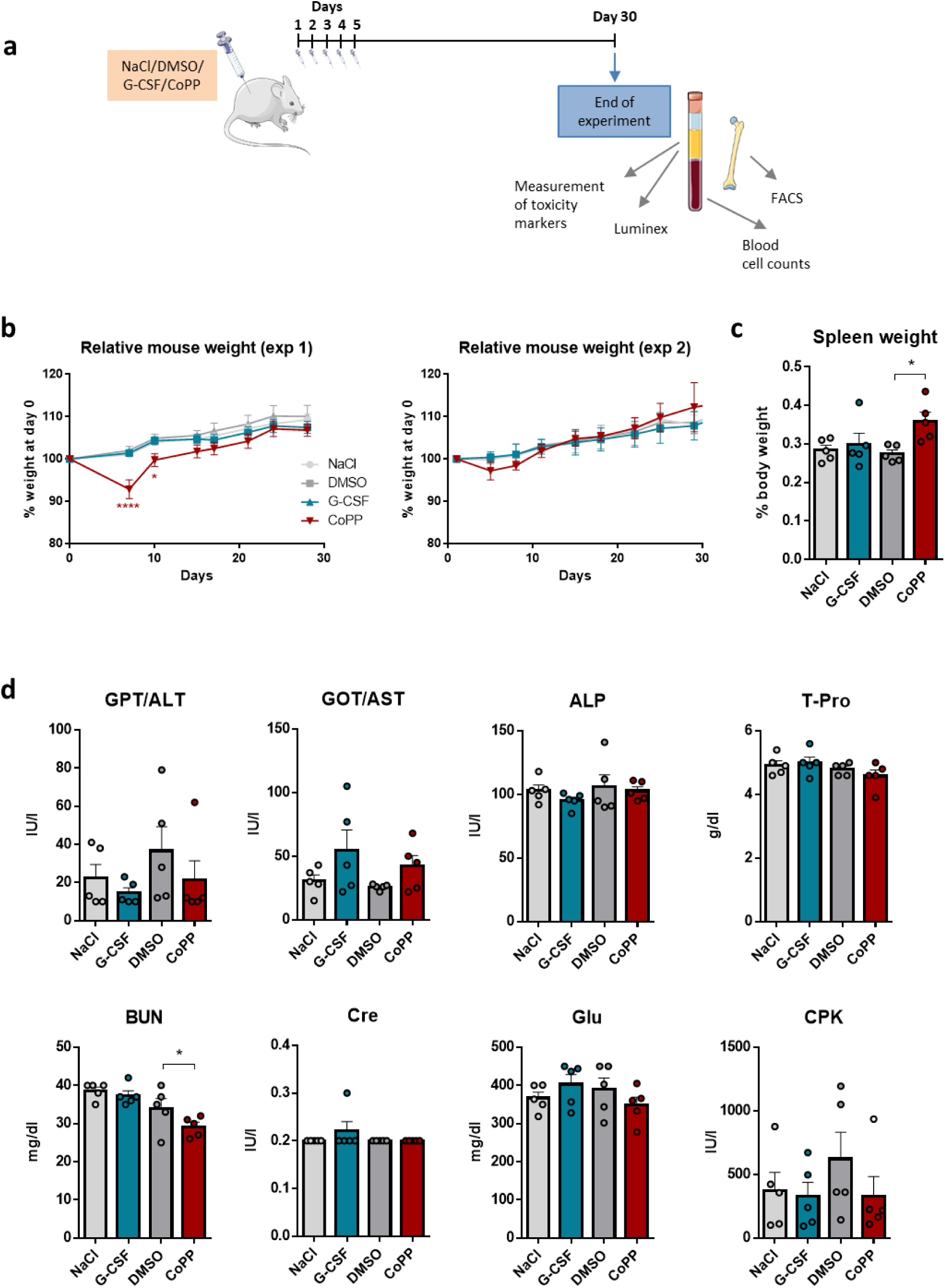
Most CoPP-induced changes resolve within 25 days after stopping the mobilization treatment. **a** Experimental scheme. C57BL/6J mice were injected daily with 10 mg/kg CoPP, 250 µg/kg recombinant G-CSF, or solvent controls (NaCl, DMSO) for 5 days. Samples were collected 25 days after the final injection. The experiment was repeated twice. **b** Relative mouse weight (data shown separately for the two independent experiments). CoPP causes a transient decrease in weight, with mice returning to their initial weight 5 days after the end of treatment. **c** Relative spleen weight. Mice treated with CoPP show an increase in spleen weight 25 days after treatment. **d** Organ function biomarkers in plasma, measured using the SpotChem EZ Chemistry Analyzer. Most liver and kidney function markers are similar between mobilized and non-mobilized mice 25 days after treatment, with the exception of blood urea nitrogen (BUN), which remains significantly lower in CoPP-treated mice (GLP/ALT – alanine transaminase, GOT/AST – aspartate aminotransferase, ALP – alkaline phosphatase, T-Pro – total protein, Cre – creatinine, BUN – blood urea nitrogen, glucose, CPK – creatine phosphokinase). Data information: Results are shown as mean + SEM; B: Two-way ANOVA with Tukey’s post-hoc test, C-D: One-way ANOVA with Sidak’s post-hoc test; n = 5 mice per group; *p ≤ 0.05; **p ≤ 0.01; ***p ≤ 0.001; ****p ≤ 0.0001

It is well-established that G-CSF treatment increases spleen weight during mobilization, and we found that CoPP had a similar effect [11]. To assess whether the spleen returned to its normal size after treatment, we measured spleen weight 25 days after CoPP treatment was terminated. We found that spleen weight remained elevated in CoPP-treated mice (0.36 ± 0.02% of total body weight) compared to DMSO-treated controls (0.27 ± 0.01%), while G-CSF-treated mice showed no such prolonged effect (0.3 ± 0.03%, Fig. 3c).

At the end of the experiment, we analyzed plasma samples for organ damage markers. One notable finding was that, unlike G-CSF, CoPP treatment led to a sustained reduction in BUN. Even 25 days post-treatment, BUN levels remained significantly lower in CoPP-treated mice (29.2 ± 1.16 mg/dl) compared to the control groups (34 ± 2.53 mg/dl for DMSO and 37.4 ± 1.2 mg/dl for G-CSF, Fig. 3d). Glucose levels, which had been reduced during 5-day CoPP treatment (Fig. 2c), returned to normal by day 30 (348.4 ± 21 mg/dl) (Fig. 3D). Other parameters, including ALT, aspartate aminotransferase (GOT/AST), ALP, T-Pro, creatinine, and creatine phosphokinase (CPK), showed no significant differences between groups, although the mean AST activity was slightly higher in the mobilized groups (both G-CSF and CoPP).

As mobilization impacts hematopoietic cell populations in both the peripheral blood and bone marrow, we evaluated its long-term effects on the hematopoietic system. Blood cell counts remained similar across all groups 25 days post treatment (Fig. 4a). The mean WBC counts ranged from 12 190 ± 1 105 cells/µl (NaCl) to 14 230 ± 1 241 cells/µl (CoPP). The proportions of leukocyte types were also consistent, with granulocyte percentages between 20.48 ± 2.1% (NaCl) and 23.67 ± 2.1% (G-CSF), monocyte percentages between 5.22 ± 0.14% (NaCl) and 6.13 ± 0.21% (CoPP), and lymphocyte percentages between 71.1 ± 2.1% (G-CSF) and 74.3 ± 2.3% (NaCl). Red blood cell parameters were largely unaffected (Fig. 4b). Hematocrit values ranged between 51 ± 1.8% in CoPP-treated mice and 55.32 ± 1.3% in the G-CSF group, hemoglobin concentrations ranged between 13.88 ± 0.46 g/dl (CoPP) and 15.06 ± 0.33 g/dl (G-CSF), with RBC counts between 10.62 ± 0.33 million cells/µl (CoPP) and 11.06 ± 0.25 million cells/µl (NaCl).

**Fig 4.**
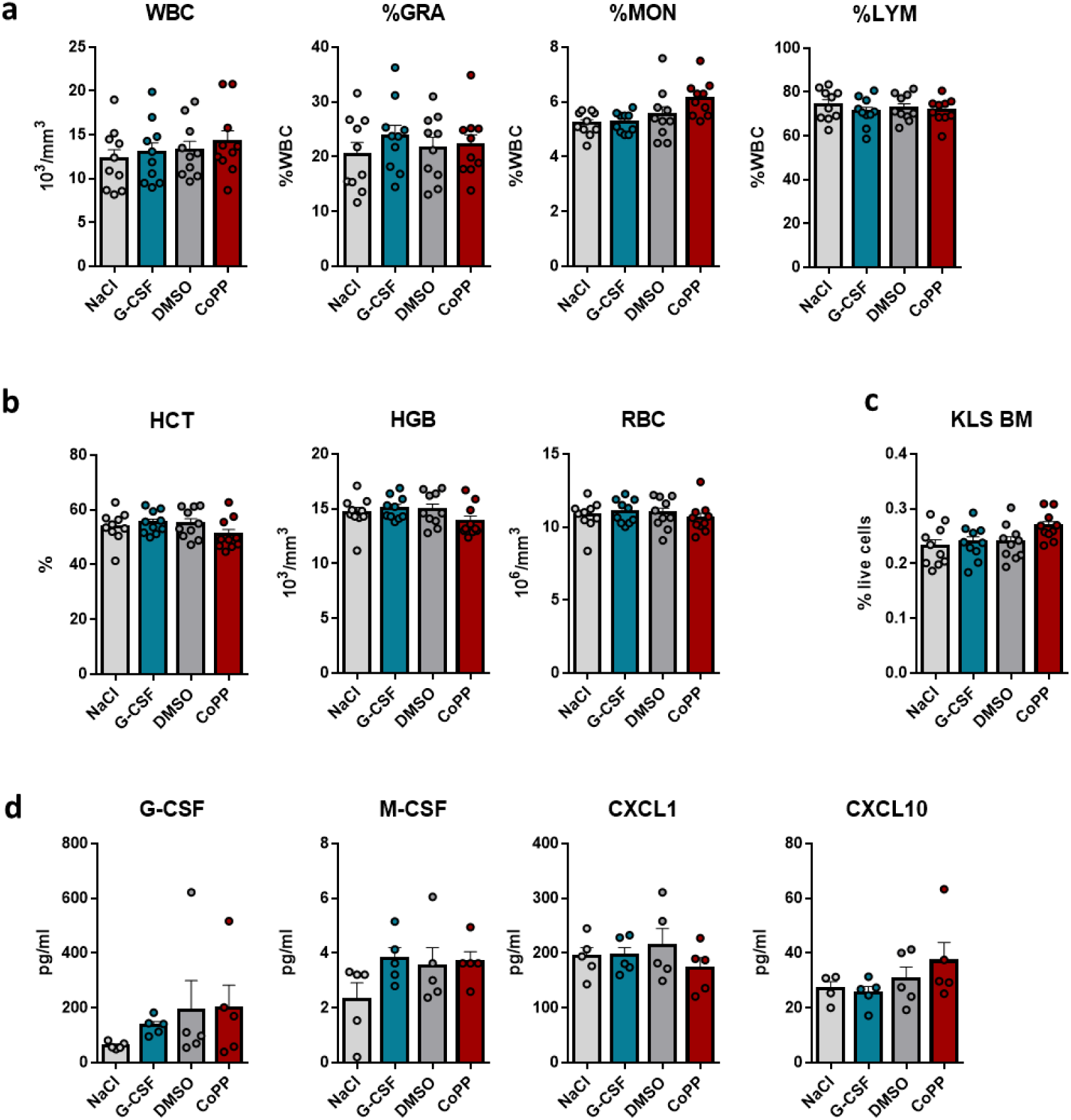
Hematological parameters and cytokine concentrations return to baseline within 25 days after the cessation of mobilization treatment. C57BL/6J mice were injected daily with 10 mg/kg CoPP, 250 µg/kg recombinant G-CSF, or solvent controls (NaCl, DMSO) for 5 days. Samples were collected 25 days after the final injection. **a** Total leukocyte count in blood. White blood cell (WBC) counts and the percentages of major leukocyte populations (%GRA, %MON, %LYM) are comparable across all groups. **b** Red blood cell parameters (HCT – hematocrit, RBC – red blood cell count, HGB – hemoglobin) are similar between groups and fall within the normal range. **c** The number of c-kit+Lineage-Sca-1+ cells in the bone marrow is comparable between mobilized (G-CSF, CoPP) and non-mobilized (NaCl, DMSO) mice. **d** Selected cytokine and growth factor concentrations in plasma, measured by Luminex assay. Cytokine concentrations induced by CoPP return to control levels 25 days after the end of mobilization treatment. Data information: Results are shown as mean + SEM; One-way ANOVA with Sidak’s post-hoc test; *p ≤ 0.05; **p ≤ 0.01; ***p ≤ 0.001; ****p ≤ 0.0001. A-C: Results were pooled from two independent experiments (n = 5 mice per group each), D: n = 5 mice per group

We previously demonstrated that 5 days of treatment with CoPP, similar to G-CSF, increases the number of various subpopulations of hematopoietic stem and progenitor cells in the bone marrow [11], which are included in the KLS phenotype (c-kit^+^ lineage^-^ Sca-1^+^) [31]. Importantly, KLS levels in the bone marrow of mice mobilized with G-CSF and CoPP returned to baseline by the end of the experiment, indicating that the hematopoietic system had regained homeostasis after 25 days (NaCl: 0.23 ± 0.01%, G-CSF: 0.24 ± 0.01%, DMSO: 0.24 ± 0.01%, CoPP: 0.27 ± 0.01% of BM live cells, Fig. 4c).

Finally, we evaluated whether the concentrations of cytokines and growth factors induced by CoPP treatment had returned to baseline by day 30. IL5, IL6, and CCL2 levels were below detection limits, and G-CSF concentrations ranged from 62.91 ± 5.8 pg/ml (NaCl) to 197.7 ± 85.6 pg/ml (CoPP), which were similar to levels typically seen in untreated or solvent-treated mice (Fig. 4d). No differences were observed in M-CSF, CXCL1, or CXCL10 levels between the groups.

In summary, most of the cellular and biochemical changes induced by five days of CoPP treatment resolve after one month.

## Discussion

In this study, we aimed to evaluate the potential of cobalt protoporphyrin (CoPP) as a novel agent for mobilizing hematopoietic stem cells (HSCs) from the bone marrow into peripheral blood. While current mobilization agents, such as granulocyte colony-stimulating factor (G-CSF), have been lifesaving by facilitating stem cell transplants, their efficacy is not universal. Some patients respond poorly, necessitating the development of new therapeutic options. Given G-CSF’s limitations, exploring alternatives like CoPP with potentially different mechanisms or improved efficacy is essential. Our previous findings showed that CoPP could mobilize cells more efficiently than G-CSF [11]. However, critical questions remained regarding its optimal dosing strategy and safety profile. Specifically, it was unclear what the minimal effective dose of CoPP was and whether this mobilization would have any long-term detrimental effects on hematopoietic system homeostasis.

We aimed to address these gaps by determining the minimal effective dose and optimal administration scheme to maximize CoPP’s mobilization potential. Furthermore, we evaluated the impact of this dose on overall safety, with a focus on potential disruptions to hematopoietic homeostasis and long-term consequences. Since successful mobilization strategies can significantly alter the bone marrow microenvironment and hematopoiesis [13,32], it was essential to assess any possible adverse effects related to these changes.

In our previous studies we used the dose of 10 mg/kg, which was effective in inducing G-CSF expression and mobilization of cells form the bone marrow to the blood [11]. However, as lower doses, such as 1 and 5 mg/kg were shown to be effective [29,33], we decided to investigate the effects of these lower doses to determine if they can similarly induce mobilization. This approach is particularly relevant given that lower doses of other metalloporphyrins have been used in clinical trials. For example, in studies by Martinez et al. [34] and Kappas et al. [35], tin mesoporphyrin (SnMP) was used to prevent or treat hyperbilirubinemia in newborns. A single dose of 6 μmol/kg of birth weight was administered, corresponding to approximately 4.53 mg/kg. More recently, a phase-2 clinical trial tested RBT-1, a combination drug of tin protoporphyrin (SnPP) and iron sucrose (FeS), in patients undergoing coronary artery bypass graft or heart valve surgery. In this trial, patients received a single intravenous infusion of either high-dose RBT-1 (90 mg SnPP/240 mg FeS) or low-dose RBT-1 (45 mg SnPP/240 mg FeS) [36].

By treating mice with 1, 5, and 10 mg/kg of CoPP, we demonstrated that CoPP induces cytokine production and mobilization in a dose-dependent manner. In addition to increasing HO-1 expression dose-dependently [37,38], CoPP has also been shown to inhibit nitric oxide synthase in the rat hypothalamus [39] and attenuate renal fibrosis in a rat model of obstructive nephropathy [37]. Interestingly CoPP dose-dependently inhibits replication of certain viruses, for example EqHV-8 in murine alveolar macrophage MH-S cells [40] or SARS-CoV-2 replication in the human lung epithelial Calu3 cells [41]. In our study, the lowest dose (1 mg/kg) induced some, but not all, of the cytokines detected after higher doses of CoPP and led to a slight increase in blood granulocytes. Although this dose was insufficient for therapeutic or experimental mobilization, it’s important to note that even this low dose affected bone marrow and blood leukocytes, which may have implications for research involving tumors, inflammation, or infections.

The effect of CoPP on weight loss is well-documented, with its mechanism studied extensively in mice [33], rats [42,43], and dogs [44]. A study by Csongradi et al. on obese melanocortin-4 receptor-deficient mice demonstrated that CoPP increases oxygen consumption, activity, and heat production, while also lowering body weight [33]. Similarly, a recent study by Rubio-Atonal et al. in rats showed that CoPP reduces food intake, water consumption, and body weight [43]. Consistent with published data, we observed a transient reduction in mouse body weight, which resolved within five days after treatment cessation. In contrast, we did not observe any weight loss in mice treated with recombinant G-CSF, so this effect is most probably not related to mobilization itself. Similarly, only in the mice treated with CoPP, we saw a transiently decreased concentration of glucose.

We also explored how CoPP affects other metabolic parameters, particularly liver and kidney markers, to better understand its broader physiological effects and potential toxicity. Since mobilization itself may elicit some side effects, we included a group of mice treated with G-CSF to observe the potential impacts of G-CSF and G-CSF-induced mobilization on the measured parameters. Interestingly, we observed decreased alkaline phosphatase (ALP) activity in the plasma of mice treated with CoPP and G-CSF. A thorough literature search did not reveal any studies reporting decreased ALP activity following mobilization or G-CSF or CoPP treatment. In contrast, several reports indicate increased ALP activity. For instance, rats treated with 100 or 300 µg/kg daily for 13 weeks exhibited increased serum ALP levels, which returned to control levels after the treatment was discontinued [45]. Additionally, a study in cancer patients found that treatment with rhG-CSF raised ALP levels, which returned to baseline 5-7 days after the completion of treatment [46].

Treatment with CoPP resulted in a significant decrease in blood urea nitrogen (BUN) levels in the plasma, an effect that was evident as early as the second day of treatment and persisted for at least 25 days after discontinuation. BUN, along with creatinine, is commonly used as a marker for acute kidney injury [47]. However, in cases of kidney injury, BUN levels typically increase rather than decrease. The absence of an increase in BUN, coupled with unchanged creatinine levels, suggests that CoPP does not impair kidney function. This is further supported by the fact that creatinine is less influenced by factors such as food intake and hydration status, making it a more reliable indicator of kidney health in this context [48]. The reduction in BUN levels observed in CoPP-treated mice may instead be linked to other mechanisms. One possibility is malnutrition, as decreased food intake has been observed in CoPP-treated mice and could explain the reduction in BUN [49]. Another potential explanation for the decreased BUN levels, along with reductions in alkaline phosphatase (ALP) activity, is the impact of CoPP on liver function. Decreased BUN levels are sometimes associated with liver dysfunction, potentially due to CoPP-induced depletion of cytochrome P450 enzymes, which are critical for hepatic metabolic processes [23]. These findings highlight the need for further studies to clarify the underlying mechanisms of BUN reduction and to assess its implications for clinical safety, particularly regarding potential effects on liver function.

By examining hematological parameters in mice treated with CoPP for 5 days and then 25 days after the treatment ended, we demonstrated that the effects of CoPP on leukocyte counts, red blood cell parameters, and hematopoietic stem and progenitor cells in the bone marrow are transient. Given the proposed therapeutic uses of CoPP [26,27,50], understanding its impact on the hematopoietic system and ensuring it does not compromise long-term hematopoiesis is essential.

In summary, this study defined the optimal dose and duration of CoPP treatment to effectively induce the mobilization of granulocytes into peripheral blood in mice. We also demonstrated that most (though not all) of the changes elicited by CoPP resolve within 30 days. These findings suggest that CoPP could potentially be used as a drug for inducing granulocyte mobilization. However, further studies are needed to elucidate the mechanisms by which CoPP affects parameters such as blood urea nitrogen and alkaline phosphatase. Additionally, if CoPP is considered for potential therapeutic use beyond mobilization, its effects on the hematopoietic system must be carefully evaluated. Although our study has some limitations, including the lack of human trials with CoPP and the potential for species-specific variability, there are several key points worth noting. While CoPP has not yet been tested in clinical trials, similar effects have been observed in other species, such as rats and dogs, suggesting that its biological activity may be consistent across different models. Additionally, other metalloporphyrins, such as tin mesoporphyrin (SnMP), have demonstrated similar mechanisms of action in both humans and mice. These findings underscore the potential of CoPP as a therapeutic agent, though further studies are necessary to fully establish its efficacy and safety in human populations.

## Author contributions

AS planned and supervised the study, analyzed the data, prepared the figures and wrote the manuscript; AB and KS assisted with the experimental design and data analysis; AB, PK, KK, IS, NKT, JFG, PK, RGG, KG, KM, AK, NBK, KS and AS performed the experiments; AB and KG helped with graph preparation, AJ provided critical support during the grant application process, which secured the funding necessary for this research; AB, IS, NKT, PK, KG, AJ and KS edited the manuscript. All the authors read and approved the final manuscript.

## Acknowledgments

The authors would like to acknowledge the Animal Facility employees for their help and support in performing animal experiments, Agnieszka Andrychowicz-Rog for technical support and all the lab members for help with experiments.

AI (ChatGPT) was used to assess and correct the text for clarity and grammar.

Funding: The study was supported by The National Centre for Research and Development, Poland (grant no LIDER/37/0133/L-11/19/NCBR/2020).

## Data availability

All experimental data is available upon request from the corresponding author. Upon publication of all project results, the data will be made publicly accessible through the RODBUK Cracow Open Research Data Repository (https://rodbuk.pl/).

